# Initiation of asexual reproduction by the AP2/ERF gene *GEMMIFER* in *Marchantia polymorpha*

**DOI:** 10.64898/2026.01.16.699827

**Authors:** Go Takahashi, Saori Yamaya, Facundo Romani, Ignacy Bonter, Kimitsune Ishizaki, Masaki Shimamura, Tomohiro Kiyosue, Jim Haseloff, Yuki Hirakawa

## Abstract

Plants can propagate their own clones through asexual reproduction. Genetic and hormonal factors regulating asexual reproduction have begun to be elucidated in the liverwort *Marchantia polymorpha*, which produce asexual propagules called gemmae within the gemma cups. Here, we report an AP2/ERF family gene, *GEMMIFER* (Mp*GMFR*), as a key regulator of asexual reproduction in *M. polymorpha*. Suppression of MpGMFR function using genome editing and amiRNA results in the loss of gemma and gemma cup formation. In contrast, activation of MpGMFR function using a dexamethasone inducible system promotes gemma and/or gemma cup formation depending on the induction conditions. Notably, transient strong activation of MpGMFR induced gemma initial cells at the meristem, which develop into mature gemmae capable of reproducing as new individuals after detachment. These results show that MpGMFR is necessary and sufficient for the initiation of asexual reproduction at the meristem. Mp*GMFR* expression was detected from the meristem through to the early stages of gemma development including the gemma cup floor cells. Mp*GMFR* expression precedes the expression of *GEMMA CUP-ASSOCIATED MYB1* (Mp*GCAM1*) in early gemma development, supporting the notion that Mp*GMFR* initiates asexual reproduction. We anticipate this finding to be a foundation to study the evolution of extra meristem formation in plant bodies that may have made a significant contribution to the prosperity of plants on land.

## Introduction

Plants have the remarkable ability to clone new individuals from their own bodies. This process, called asexual reproduction, occurs at various structures such as adventitious shoots, bulbils, tubers and rhizome buds^1^. Bryophytes often reproduce asexually through the dispersal of propagules called gemmae with species-specific morphology^2,3^. In the liverwort *Marchantia polymorpha*, discoid-shaped gemmae are produced in specialized cup-shaped structures called gemma cups which are periodically generated along the dorsal midrib of its thalloid body^4^. Recent studies on *M. polymorpha* are beginning to elucidate genetic and hormonal regulation of the gemma and gemma cup development^5^. An R2R3-MYB transcription factor, GEMMA CUP-ASSOCIATED MYB1 (MpGCAM1), has been identified as a gene highly expressed in gemma cups^6^. Molecular genetic analysis has shown that Mp*GCAM1* is required for the formation of gemma cups and gemmae, acting through the control of cell differentiation. Cytokinin signaling promotes the formation of gemma cups through upregulation of Mp*GCAM1* expression^7,8^. KARRIKIN INSENSITIVE2 (KAI2)-dependent signaling promotes gemma cup formation by upregulating the expression of a cytokinin biosynthesis enzyme, MpLONELY GUY (MpLOG), leading to the upregulation of Mp*GCAM1* expression^9,10^. Another R2R3-MYB transcription factor, SHOT GLASS (MpSTG), regulates the shape of gemma cup and the gemma development^11^. In addition, several genes are reported to be required for morphogenesis during early gemma development. Mp*ROOT HAIR DEFECTIVE SIX-LIKE1* (Mp*RSL1*) transcription factor gene, targeted by the microRNA *FEW RHIZOIDS1* (Mp*FRH1*), is required for cellular outgrowth at the floor of gemma cup, which will later develop into gemmae^12,13^. The single copy *RHO of Plant* (Mp*ROP*) gene and its regulatory factors are essential for the morphogenesis of various tissues and organs including gemmae and gemma cups^14–17^. Signaling of the plant hormones auxin, ethylene and jasmonate affect the morphology of gemmae^18–20^. Although these studies have identified a number of genes regulating gemma development, the key genes controlling the initiation of the gemma cell lineage remain unknown.

We have previously shown that MpCLAVATA3/EMBRYO SURROUNDING REGION-related 2 (MpCLE2) peptide signaling negatively regulates the formation of gemma cups, in addition to its function in stem cell identity in the meristem located in the apical notch^21–23^. In a transcriptome analysis, we have identified several differentially expressed transcription factor (DETF) genes in Mp*CLE2* gain-of-function transgenic lines. Among them, JINGASA (MpJIN), affects stem cell fate in the meristem by promoting periclinal cell division^24^. In this study, we report that another DETF, GEMMIFER (MpGMFR)/MpERF14, plays a key role in the initiation of asexual reproduction. This finding would provide a molecular clue to understanding how plant cell fate is regulated in asexual reproduction.

## Results

### Mp*GMFR* is essential for the formation of gemma cups and gemmae

To understand the function of Mp*GMFR*, we generated loss-of-function alleles using the CRISPR-Cas9 genome editing. Two independent frame-shift alleles (Mp*gmfr-1*^*ge*^ and Mp*gmfr-2*^*ge*^) were obtained (Fig. S1A and S1B), and both resulted in complete loss of gemma cup and gemma formation. For the quantification, we observed 2-week-old thallus developed from an explant containing an apical notch. In wild type, all examined plants formed gemma cups on the dorsal surface (Fig. 1A). In contrast, none of the Mp*gmfr-2*^*ge*^ plants formed gemma cups and gemmae (Fig. 1B and 1C). This Mp*gmfr*^*ge*^ phenotype was partially complemented by the introduction of a gRNA-resistant Mp*GMFR* expressed under its own promoter (Fig. S1C and S1D). To further elucidate the function of Mp*GMFR*, we generated estrogen-inducible artificial microRNA lines targeting Mp*GMFR* mRNA using the XVE transactivation system (_*pro*_Mp*E2F:XVE*>>*amiR-*Mp*GMFR*). In RT–qPCR assays, Mp*GMFR* mRNA levels were decreased in an estrogen-dependent manner in the _*pro*_Mp*E2F:XVE*>>*amiR-*Mp*GMFR* plants compared to wild-type plants (Fig. S1E). The _*pro*_Mp*E2F:XVE*>>*amiR-*Mp*GMFR* plants grown for three weeks on estrogen-free medium formed gemma cups, but those on estrogen-containing medium did not (Fig. 1D and 1E). We transferred the 2-week-old plants from estrogen-free to estrogen-containing medium and grown further for 1 week, or *vice versa*. The transferred plants lost the formation of gemma cups and gemmae in an estrogen-dependent, reversible manner (Fig. 1F and 1G). Collectively, these data show that Mp*GMFR* is required for the formation of gemma cups and gemmae in *M. polymorpha*.

**Fig. 1:**
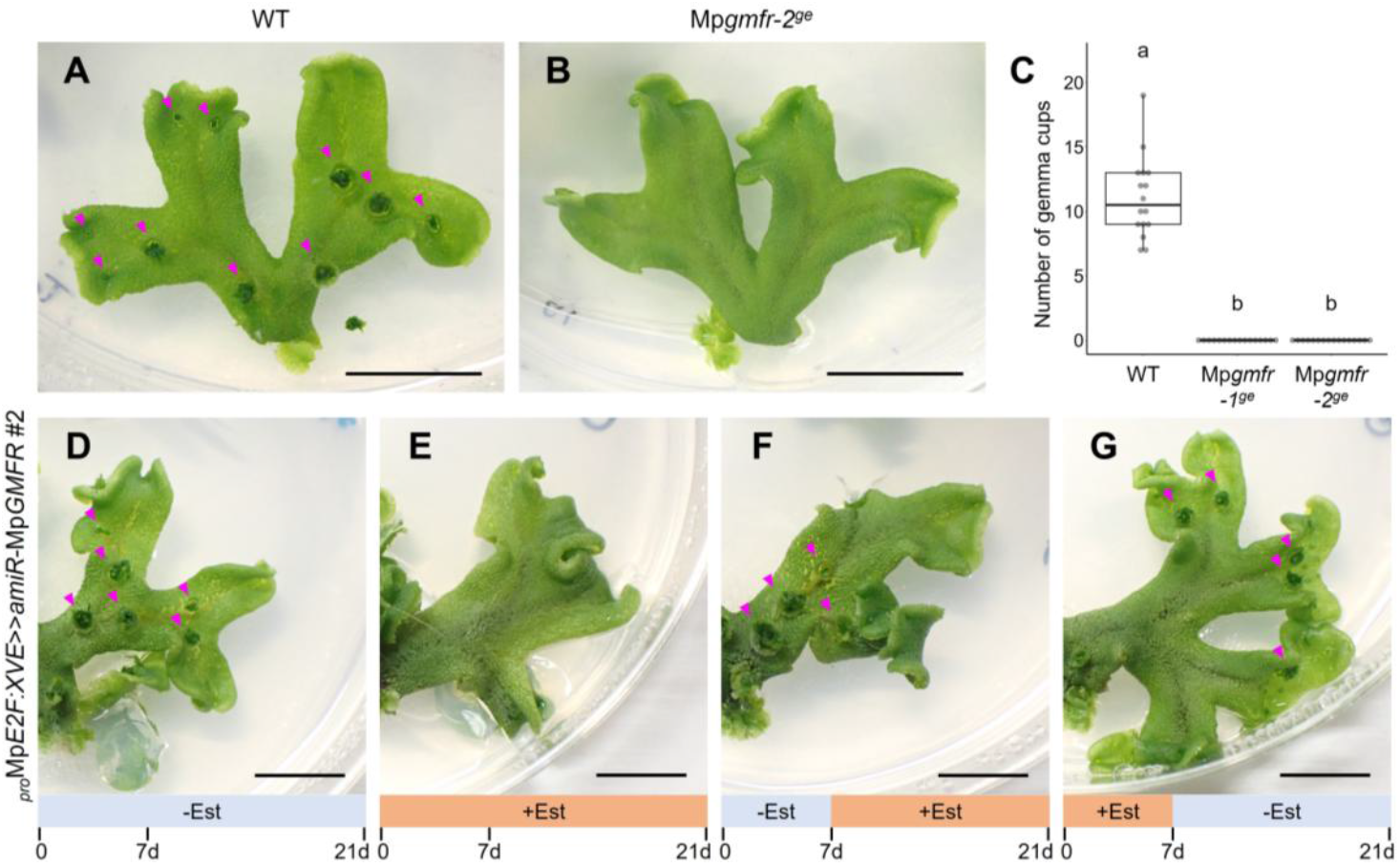
Mp*GMFR* is essential for the formation of gemma cups and gemmae. (**A and B**) Two-week-old thalli grown from the tips of thalli of wild type (**A**) and Mp*gmfr-2*^*ge*^ (**B**). (**C**) The number of gemma cups (n=16). (**D–G**) Effects of Mp*GMFR* knockdown on gemma cup formation in _*pro*_Mp*E2F:XVE>>amiR-*Mp*GMFR*. Expression of *amiR-*Mp*GMFR* was induced by β-estradiol (Est) during different periods within the 21 days as indicated below the panels. In **A, B, D–G**, arrowheads indicate gemma cups. In **C**, the boxes show the median and interquartile range, and the whiskers show the 1.5x interquartile range. Individual data points are plotted as dots. Two-way ANOVA with Tukey’s post hoc test in **C**. Means sharing the superscripts are not significantly different from each other, p < 0.05. Scale bars represent 1 cm (**A**,**B**,**D–G**).

Molecular phylogenetic analysis has shown that *M. polymorpha* possesses three genes, Mp*ERF1* (Mp1g20040), Mp*ERF14*/*GMFR* (Mp4g00380) and Mp*ERF20*/*LAXR* (Mp5g06970) in class VIII of the AP2/ERF (ERF-VIII) family^25,26^. A phylogenetic tree with ERF-VIII genes from various land plant species (*Arabidopsis thaliana, Amborella trichopoda, Ginkgo biloba, Salvinia cucullata, Azolla filiculoides, Selaginella moellendorffii, Sphagunum fallax, Physcomitrium patens, Marchantia polymorpha* and *Anthoceros agrestis*), suggests that ERF-VIII can be divided into three subgroups, each of which contains single *M. polymorpha* gene (Fig. S2A). To verify the functional redundancy among ERF-VIII genes in *M. polymopha*, we conducted complementation test of the Mp*gmfr*^*ge*^ phenotype by introducing the coding sequences of Mp*ERF1* and Mp*LAXR*, respectively, expressed under Mp*GMFR* promoter. Neither construct complemented the Mp*gmfr*^*ge*^ phenotype, suggesting that Mp*GMFR* plays an essential function in the gemma cup and gemma formation that cannot be replaced by either Mp*ERF1* or Mp*ERF20*/*LAXR* (Fig. S2B).

### Temporal activation of MpGMFR induces ectopic gemmae formation

In *M. polyrmopha*, gemmae are generated from the floor cells at the bottom of gemma cup. At the initiation of gemma development, a gemma cup floor cell protrudes from the epidermal surface and then divides transversely twice into three cells: a gemma cell, a stalk cell and a basal cell^27,28^ (Fig. 2A). While the stalk cell stops further divisions, the gemma cell continues to divide and grows into the discoid-shaped gemmae with two apical meristematic notches at bilateral sites. Thus, the gemma cell acts as the initial cell for the development of gemma.

**Fig. 2:**
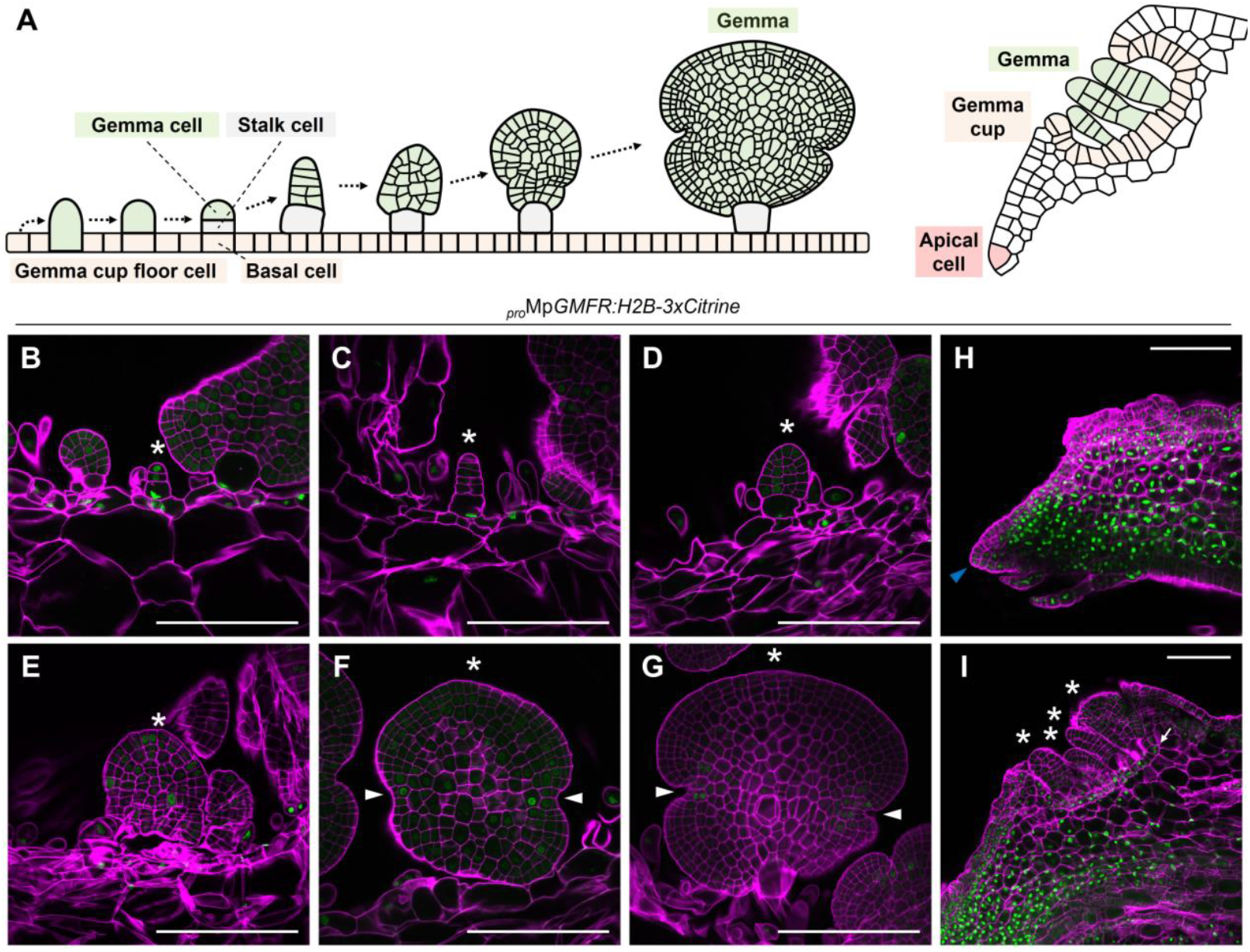
Expression patterns of Mp*GMFR* in developing gemma and gemma cup. (**A**) Schematic illustration of early development of gemma (left) and developing gemma cup (right, modified from Barnes and Lang, 1908^28^). (**B–I**) Confocal imaging of _*pro*_Mp*GMFR:H2B-3xCitrine* in developing gemmae (**B–G**) and cross-sections of thallus (**H**,**I**). Cell walls were stained with SCRI Renaissance 2200 (SR2200). Asterisks show developing gemmae. Blue and white arrowheads indicate a (sub)apical cell and apical notches, respectively. An arrow indicates the layer of gemma cup floor cells. Scale bars represent 100 μm (**B–I**).

To analyze the expression patterns of Mp*GMFR* during asexual reproduction, we observed transcriptional-fusion reporter lines of Mp*GMFR* (_*pro*_Mp*GMFR:H2B-3xCitrine*) under confocal laser scanning microscopy in the cross-section of gemma cup cleared with the iTOMEI method^29^. The fluorescence signals of _*pro*_Mp*GMFR:H2B-3xCitrine* were detected in the gemma cup floor cells. (Fig. 2B–G). In the initial stage of gemma development, the signals were also detected in both the gemma cells and the stalk cell (Fig. 2B), but were lost after the subsequent several divisions of the gemma cell (Fig. 2C). In a later stage, the signals were scattered within the developing gemmae (Fig. 2D and 2E). After the formation of meristematic notches at bilateral sites of the gemmae, the signals were restricted to cells near the notches (Fig. 2F and 2G). This pattern is consistent with the promoter activity observed in gemmae (https://mpexpatdb.org/)^30^. In the vertical longitudinal section of the apical notch of mature plants, the signals were broadly detected at the meristem, except for the central part near the apical cells (Fig. 2H). The signals were detected the floor cells of immature gemma cup but was not in gemmae developing in the cup (Fig. 2I). These data suggest that Mp*GMFR* expression is associated with meristematic region, gemma cup floor cells and the very early stages of gemma development.

To understand the function of Mp*GMFR*, we first generated overexpression lines using constitutive Mp*EF1α* promoter. The _*pro*_Mp*EF1α:*Mp*GMFR* plants result in ball-shaped structure composed of small immature thalli (Fig. 3A). To analyze the short-term effects of overexpression, we generated inducible overexpression lines (_*pro*_Mp*EF1α:*Mp*GMFR-GR*) in which MpGMFR proteins fused with glucocorticoid receptor (GR) are expected to be translocated into the nucleus by dexamethasone (DEX) treatment. In DEX-containing medium, 4-day-old _*pro*_Mp*EF1α:*Mp*GMFR-GR* plants formed protruding cells around the apical notch (Fig. 3B). These cells can be stained by _*pro*_Mp*YUC2:GUS* and have morphology similar to the gemma cell in the initial stage of gemma development. However, 9-day-old _*pro*_Mp*EF1α:*Mp*GMFR-GR* plants grown on DEX medium did not form gemmae and only formed ball-shaped structures composed of immature small thalli, phenocopying the constitutive overexpression lines (Fig. S3). Since Mp*GMFR* expression is suggested to be transiently active in early development of gemmae (Fig. 2A and 2B), we hypothesized that the continuous DEX induction of MpGMFR compromised normal development of gemmae. To avoid this, 4-day-old _*pro*_Mp*EF1α:*Mp*GMFR-GR* plants grown on DEX-containing medium were transferred to DEX-free medium for further growth. Five days after the transfer, a number of gemmae were formed on both dorsal and ventral sides of thallus (Fig. 3C). The induced gemmae can be detached with water and contain apical notches for further growth (Fig. 3D). These notches contained apical and subapical cells like the stem cell zone in wild-type plants (Fig. 3E). In contrast to wild-type gemmae that have two apical notches at bilateral sites, the number of apical notches varied between one to four among the individual induced gemmae (Fig. 3F). In addition to the apical notches, the induced gemmae had a trace of stalk, with the cell arrangement similar to those in wild type (Fig. 3E). Although these gemma were smaller than those of wild type in the overall size (ground cover area), they can grow normally on the medium (Fig. 3D and 3G). Consistently, 5-day-old _*pro*_*35S*:Mp*GMFR-mVenus* gemmalings formed ectopic gemma on the surface (Fig. 3H). Activity of other ERF-VIII genes, Mp*ERF1* and Mp*LAXR*, were examined by generating _*pro*_Mp*EF1α:*Mp*ERF1-GR* and _*pro*_Mp*EF1α:*Mp*LAXR-GR* lines. In both cases, gemmae were not induced unlike _*pro*_Mp*EF1α:*Mp*GMFR-GR* lines, supporting the notion that ERF-VIII genes do not exert redundant function in *M. polymorpha* (Fig. S4). Collectively, these data show that temporal induction of MpGMFR leads to the ectopic formation of gemmae capable of reproducing as new individuals.

**Fig. 3:**
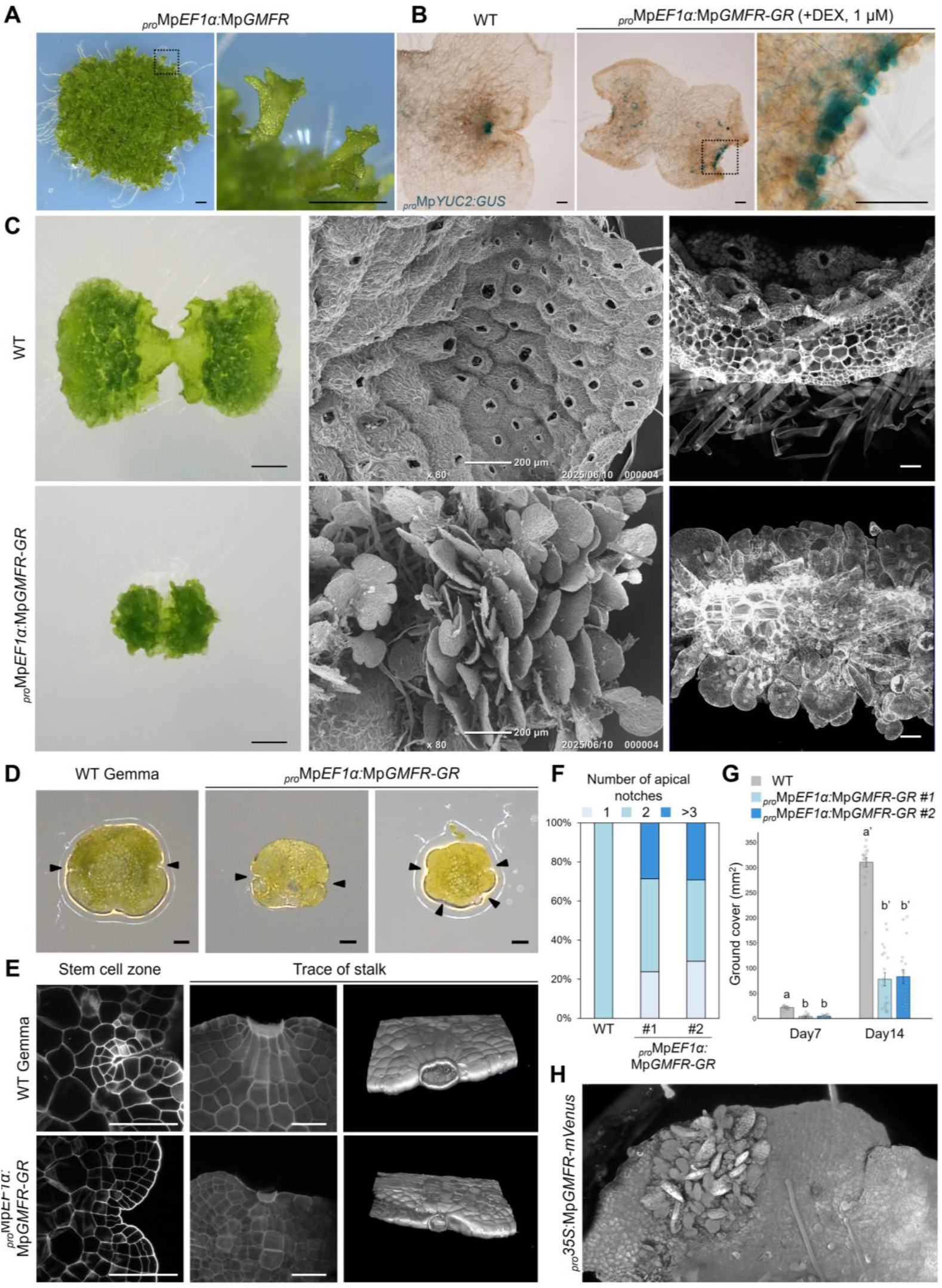
Temporal activation of MpGMFR induces ectopic gemmae formation. (**A**) Morphology of _*pro*_Mp*EF1α:*Mp*GMFR* plant. The right panel shows the magnified image of a dashed line square in the left panel. (**B**) Morphology of apical notch with _*pro*_Mp*YUC2:*GUS marker in 4-day-old plants grown from gemmae. The right panel shows the magnified image of a dashed line square in the center panel. (**C**) Morphology of 9-day-old plants grown from gemmae. Wild-type plants were grown for 9 days on normal medium. _*pro*_Mp*EF1α:*Mp*GMFR-GR* plants were first grown on 1 µM DEX-containing medium for 4 days, and transferred to DEX-free medium and grown for 5 days. Overall morphology (left), SEM images of the surface of thallus (center) and 3D-reconstructed images of the cross-section of the thallus in confocal imaging (right) were shown. (**D**) Morphology of gemmae of wild type (left) and _*pro*_Mp*EF1α:*Mp*GMFR-GR* (right, two panels). Arrowheads show notches. (**E**) Confocal imaging of the stem cell zone and a trace of stalk in gemmae of wild type (top) and _*pro*_Mp*EF1α:*Mp*GMFR-GR* (bottom). The traces of stalk were shown by 3D-reconstructed images. (**F**) The frequency of gemmae with different numbers of apical notches (n=20–23). (**G**) Ground cover area of 7 and 14-day-old plants grown from gemmae (n=18–20). (**H**) 3D-reconstructed images of 5-day-old _*pro*_*35S:*Mp*GMFR-mVenus* gemmaling. In **F**, data are represented by mean (bars) and individual data points (dots). Two-way ANOVA with Tukey’s post hoc test in **F**. Means sharing the superscripts are not significantly different from each other, p < 0.05. In **C** and **E**, cell walls were stained with SR2200. Scale bars represent 1 mm (**A** and **C**, left), 200 μm (**C**, center), 100 μm (**B, C**, right and **D**) and 50 µm (**G**).

To investigate the timing of gemma initiation after MpGMFR induction, we observed apical notches of _*pro*_Mp*EF1α:*Mp*GMFR-GR* plants grown for 1 to 4 days on 1 µM DEX-containing medium. At day 2, protruding cells that have undergone a transverse division was observed at the apical notch (Fig. 4A). The protruding cells divided transversely again to produce a gemma cell and a stalk cell at day 3. At day 4, the gemma cells divided further while the stalk cell remained inactive in cell division, recapitulating the division patterns in early development of wild-type gemmae. In some cases, the cell division patterns of the gemma cell were unusual compared to wild type, which might be the cause of the malformed gemmae (Fig. 3D and 3F). In the 3D-reconstructed images of apical notch area of the parent plants, the induced gemmae can be observed on both dorsal and ventral surfaces in _*pro*_Mp*EF1α:*Mp*GMFR-GR* plants while no gemmae were observed in wild type (Fig. 4B and S5). These data indicate that MpGMFR controls the initiation of gemma cell lineage in the meristem.

**Fig. 4:**
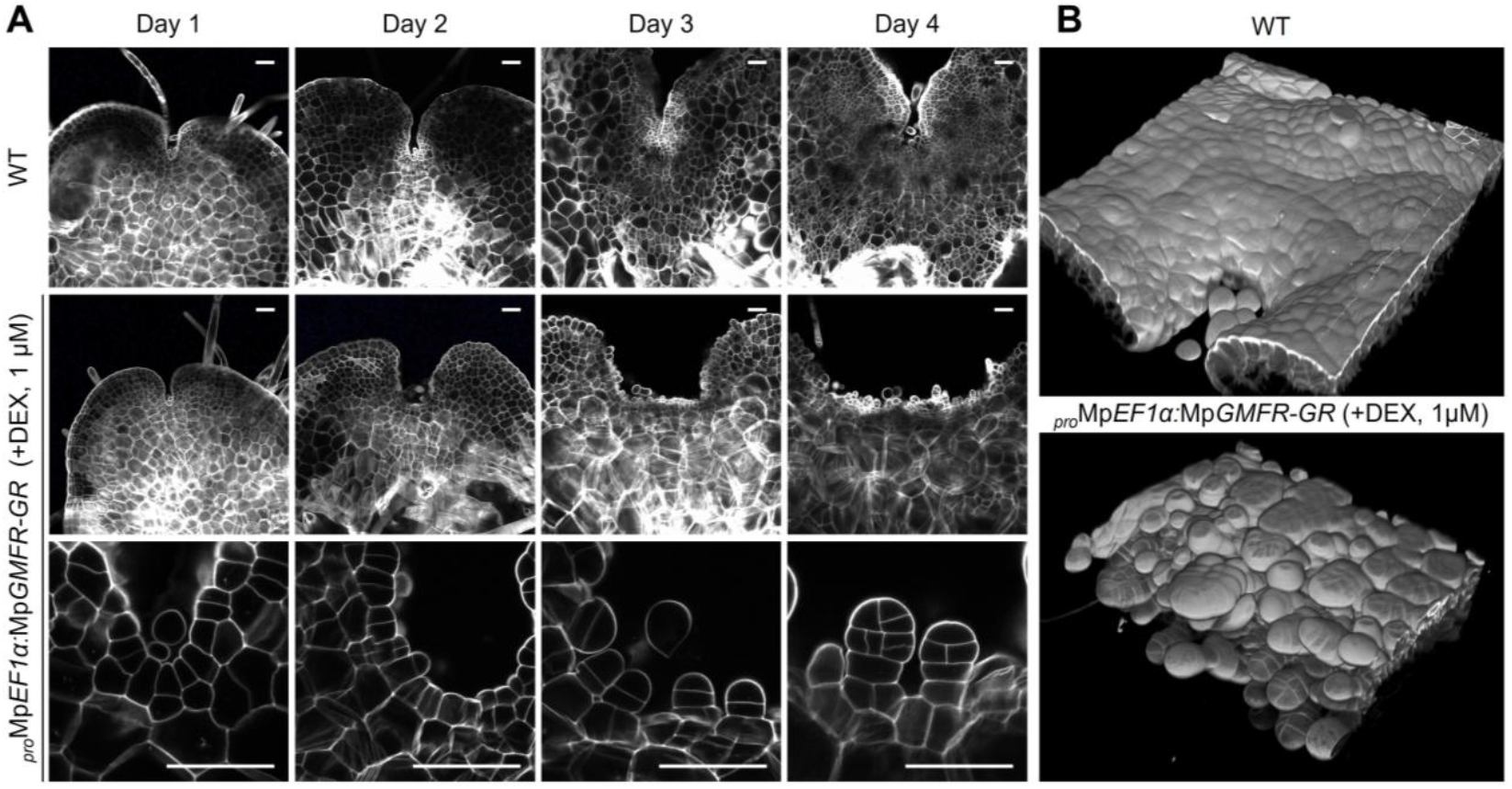
MpGMFR initiates the gemma cell lineage in the meristem. (**A**) Confocal images of the apical notch in wild type and _*pro*_Mp*EF1α:*Mp*GMFR-GR*. Panels in the bottom row show magnified images of _*pro*_Mp*EF1α:*Mp*GMFR-GR* in the middle row. (**B**) 3D-reconstructed apical notches in 4-day-old plants of wild type and _*pro*_Mp*EF1α:*Mp*GMFR-GR*. Cell walls were stained with SR2200. Scale bars represent 50 μm (**A**).

### Weak induction of MpGMFR promotes gemma cup formation

To examine the concentration-dependent effects of MpGMFR, _*pro*_Mp*EF1α:*Mp*GMFR-GR* plants were grown on growth media containing different concentrations of DEX (Fig. S6). In contrast to the 1 µM DEX treatment that only induced gemma-like tissues, 10 nM DEX treatment induced the formation of semicircular gemma cups near the apical notches, which contains gemma-like tissues inside, in 7-day-old plants (Fig. 5A). The rim structure of the induced gemma cups in the Mp*GMFR-GR* lines were similar to that of wild-type gemma cups (Fig. 5B). These data suggest that MpGMFR can also induce the formation of gemma cups.

**Fig. 5:**
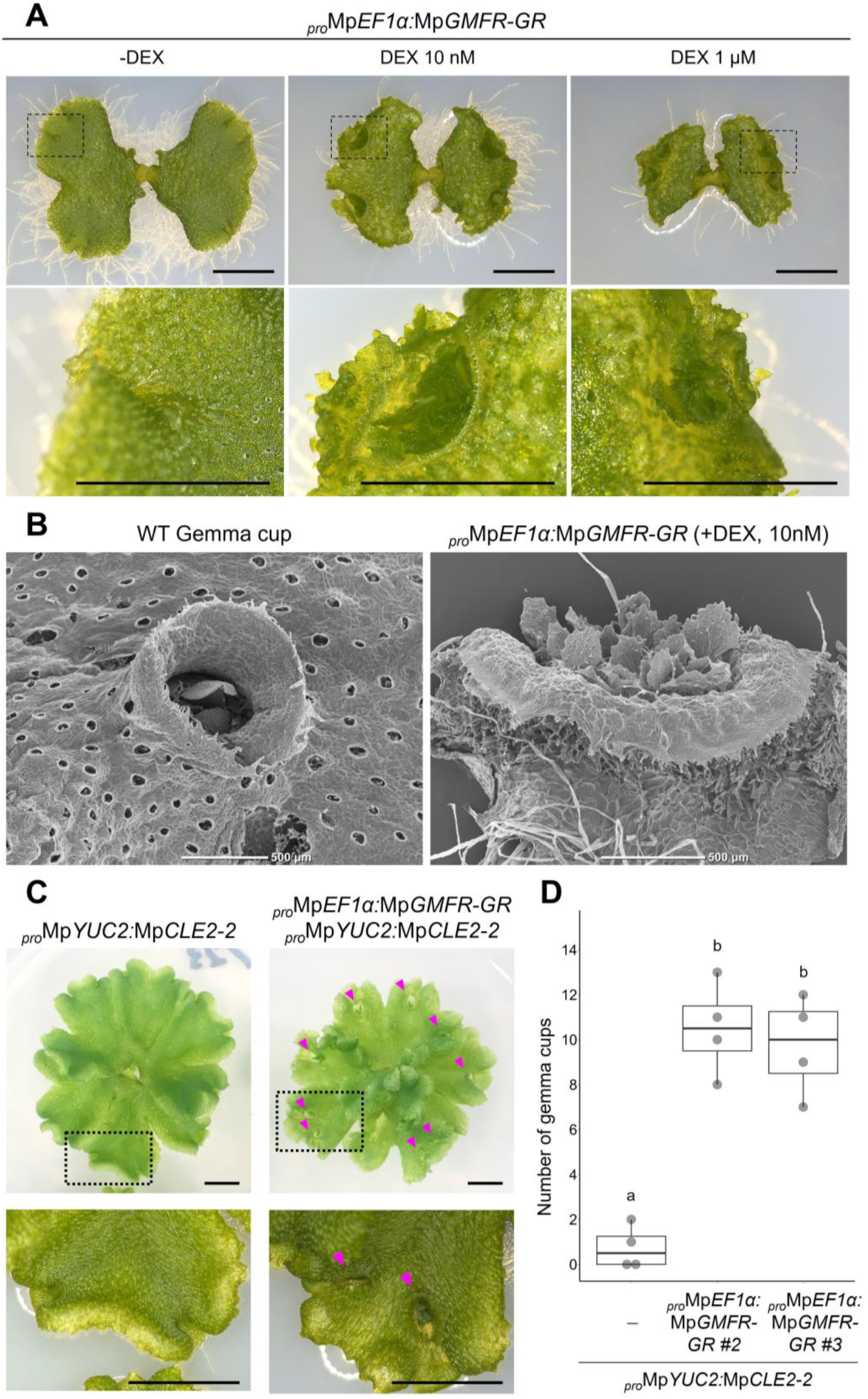
Weak induction of MpGMFR promotes gemma cup formation. (**A**) Effects of different concentrations of DEX on gemma cup formation in _*pro*_Mp*EF1α:*Mp*GMFR-GR* plants grown for 7 days from gemmae. Panels in the bottom row show the magnified images of dashed line squares in the panels above. (**B**) SEM images of the gemma cups observed in 20-day-old wild-type plant and 7-day-old _*pro*_Mp*EF1α:*Mp*GMFR-GR* plant. (**C**) Effects of MpGMFR induction in Mp*CLE2* overexpression plant. Twelve-day-old plants grown on normal medium were transferred to 10 nM DEX-containing medium and further grown for 8 days. Panels in the bottom row show the magnified images of dashed line squares in the panels above. (**D**) The number of gemma cups (n=4). In **D**, the boxes show the median and interquartile range, and the whiskers show the 1.5x interquartile range. Individual data points are plotted as dots. Two-way ANOVA with Tukey’s post hoc test in **D**. Means sharing the superscripts are not significantly different from each other, p < 0.05. Scale bars represent 5 mm (**A, C**) or 500 μm (**B**).

To examine the functional relationship between Mp*CLE2* and Mp*GMFR* genes, we generated _*pro*_Mp*EF1α:*Mp*GMFR-GR* lines in the background of a gain-of-function allele of Mp*CLE2*, _*pro*_Mp*YUC2:*Mp*CLE2-2*^21^. In this background, overexpression of Mp*CLE2* results in reduced gemma cup formation and decreased Mp*GMFR* expression levels^22,24^. In the experiment, gemmae were first grown on DEX-free medium for 12 days, followed by an additional 8 days on 10 nM DEX-containing medium. As a result, two independent Mp*GMFR-GR* lines showed a significant increase in the number of gemma cups compared to the background line (Fig. 5C and 5D), suggesting that Mp*CLE2* overexpression phenotypes on gemma cup formation can be suppressed by restoring the reduced Mp*GMFR* expression.

### Genetic interaction of Mp*GMFR* and Mp*GCAM1*

Loss-of-function alleles of Mp*GCAM1* are deficient in the gemma cup and the gemma^6^, similar to those of Mp*GMFR*. To analyze the functional interaction between these genes, we introduced frame-shift mutations in Mp*GCAM1* by CRISPR-Cas9 genome editing in the _*pro*_Mp*EF1α:*Mp*GMFR-GR* background (Fig. S7). As expected, gemma and gemma cup formation was completely lost in these lines, and we used one of them for further analysis. To analyze the effects on gemma cup formation, explants containing an apical notch were grown for 7 days on 1 µM DEX-containing medium, followed by an additional 7 days on DEX-free medium. Under these conditions, gemma-like tissues were formed in _*pro*_Mp*EF1α:*Mp*GMFR-GR* plants. In contrast, such gemma-like tissues were not observed in the Mp*gcam1*^*ge*^ _*pro*_Mp*EF1α:*Mp*GMFR-GR* line (Fig. 6A). We also could not observe protruding cells or early-stage gemmae in this line, suggesting that both Mp*GMFR* and Mp*GCAM1* are essential for the initiation of gemma development.

**Fig. 6:**
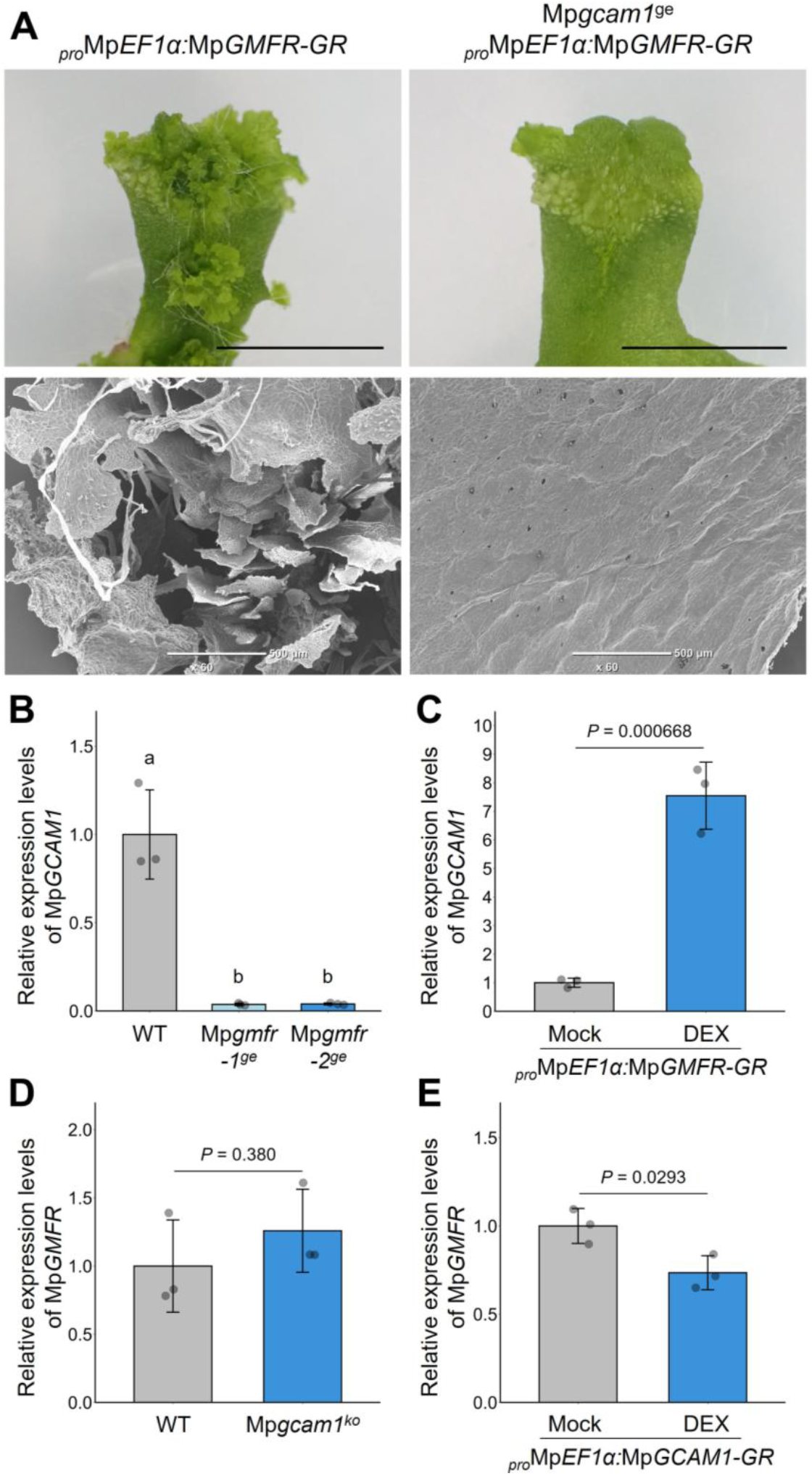
Genetic interaction of Mp*GMFR* and Mp*GCAM1*. (**A**) Effect of MpGMFR induction in Mp*gcam1*^*ge*^ plant. Seven-day-old plants grown on 1 μM DEX-containing medium were transferred to DEX-free medium and further grown for 7 days. Morphology of apical notch (top). SEM images of the surface of plants (bottom). (**B–C**) Relative expression levels of Mp*GMFR*. Six-day-old thalli of the WT and Mp*gcam1*^*ko*^ plants (**B**). Four-day-old gemmalings of _*pro*_Mp*EF1α:*Mp*GCAM1-GR* on mock or 1 µM DEX-containing medium (**C**). (**D–E**) Relative expression levels of Mp*GCAM1*. Six-day-old thalli (**D**) and four-day-old gemmalings (**E**) are examined. In **B–E**, the expression levels were normalized by Mp*APT* expression. In **B–E**, data are represented by mean (bars) and individual data points (dots). Two-way ANOVA with Tukey’s post hoc test in **B** and Student’s t test in **C–E**. Means sharing the superscripts are not significantly different from each other, p < 0.05. In **C**,**D and E**, *P*-value is indicated above each pair of bars. Scale bars represent 1 cm (**A**) or 500 μm (**B**).

To further analyze the relationship, we compared expression patterns of these genes during gemma development. While the Mp*GMFR* expression was detected in the gemma cup floor cells and early development of gemmae (Fig. 2), the fluorescence signal of the Mp*GCAM1-Citrine* knock-in allele was strongly detected throughout the gemma development but was barely detectable in the gemma cup floor cells^6,11^ (Fig. S8), suggesting that Mp*GMFR* expression precedes Mp*GCAM1* expression in gemma development. We further examined the expression levels of each gene in the mutants of the other gene by RT-qPCR assays. In the 6-day-old plants grown from explants containing an apical notch, the Mp*GCAM1* mRNA level was decreased to about 4 % in Mp*gmfr*^*ge*^ lines compared to wild type (Fig. 6B). In the 4-day-old gemmalings of an Mp*GMFR-GR* overexpression line grown with DEX, the Mp*GCAM1* mRNA level was increased to about 750 % compared to mock treatment (Fig. 6C). On the other hand, in the 6-day-old plants grown from explants, the Mp*GMFR* mRNA level was not significantly affected in a Mp*gcam1*^*ko*^ line^6^ compared to wild type (Fig. 6D). In the 4-day-old gemmalings of an Mp*GCAM1-GR* overexpression line grown with DEX, the Mp*GMFR* mRNA level was decreased to about 74 % compared to mock treatment (Fig. 6E). These data suggest that Mp*GMFR* promotes Mp*GCAM1* expression directly or indirectly while Mp*GCAM1* has little impact on Mp*GMFR* expression. Collectively, these data suggest that gemma development is initiated through the stepwise function of Mp*GMFR* and Mp*GCAM1*.

## Discussion

Due to its nature of autonomous asexual reproduction, the liverwort *Marchantia polymorpha* has become an important model organism for studying the molecular mechanisms of asexual reproduction. In this study, we identified Mp*ERF14*/*GMFR* as a key regulator of the initiation of asexual reproduction. Molecular genetic analysis shows that Mp*GMFR* is required for the development of gemma cups and gemmae while overexpression of Mp*GMFR* induces gemma cups or gemmae in a dose-dependent manner. When induced weakly, MpGMFR acts as a positive regulator of gemma cup development, which is consistent with the inhibitory function of MpCLE2 in gemma cup development because the expression of Mp*GMFR* is suppressed by MpCLE2 peptide signaling. Strong short-term induction of MpGMFR activity results in the formation of ectopic gemmae with a junction to the stalk and meristematic notches, which enable gemmae to be detached from the parental plant and grow independently in a remote location. In contrast, continued overexpression of Mp*GMFR* interferes with normal development of gemmae, suggesting the importance of spatio-temporal regulation of Mp*GMFR* expression during gemma development. Consistently, the expression of Mp*GMFR* is observed in the gemma cup floor cells and the early stages of gemma development, but it is gradually restricted near the meristems during gemma development.

Previous studies show that Mp*GCAM1* is required for gemma and gemma cup formation^6,8–10^. In this study, we show that Mp*GCAM1* expression is substantially reduced in the Mp*GMFR* loss-of-function alleles while it is increased after the induction of MpGMFR. The formation of ectopic gemmae in the Mp*GMFR-GR* lines requires Mp*GCAM1*, suggesting that Mp*GCAM1* may act in the gemma cell lineage to reinforce and/or maintain the cell identity. In contrast, Mp*GMFR* expression was not significantly affected in the Mp*GCAM1* loss-of-function allele. In the expression analysis with fluorescent protein reporters, Mp*GMFR* expression slightly precedes the Mp*GCAM1* expression during asexual reproduction. Collectively, these data suggest that early stage of gemma development can be further divided into two steps, controlled by MpGMFR and MpGCAM1, respectively.

Our phylogenetic analysis suggests that class VIII of AP2/ERF family proteins can be classified into three subgroups, each comprising genes from major lineages of land plants including single *M. polymorpha* gene (Mp*ERF1*, Mp*ERF20*/*LAXR* and Mp*ERF14*/*GMFR*). Thus, three distinct ERF-VIII genes were already present in the last common ancestor of land plants while their closest homolog is unclear in the ERF family of streptophyte algae^25^. In *Arabidopsis thaliana*, At*ERF84* which belongs to the subgroup containing Mp*GMFR* has been reported as a positive regulator in drought resistance while its function in development is unclear^31^. In *M. polymorpha*, Mp*ERF20*/*LAXR* regulates cellular reprogramming during the tissue regeneration in response to the reduction of the auxin level after the decapitation of meristematic notch^32^. Mp*ERF20*/*LAXR* is a homolog of *ENHANCER OF SHOOT REGENERATION 1*/*DORNRÖSCHEN (ESR1*/*DRN)* in *A. thaliana*, which promotes stem cell formation in the shoot regeneration^33^. In the moss *P. patens*, Pp*ESR* genes promote gametophore apical cell identity whose expression is regulated by cytokinin signaling^34^, implying deep evolutionary conservation of these genes in controlling the stem cell identity in land plants. The function of Mp*ERF14*/*GMFR* in asexual reproduction has a similarity to the ESR/LAXR genes in terms of regulation of cell identity leading to the formation of new meristems even though Mp*ERF20*/*LAXR* cannot replace the activity of MpERF14/GMFR.

In summary, we have presented the identification of Mp*ERF14*/*GMFR* as a molecular trigger of the asexual reproduction in *M. polymorpha*. This work provides a foundation for future studies to elucidate how plants evolved the formation of extra meristems from their body that may have made a significant contribution to their prosperity on land.

## Acknowledgements

We thank Ikuko Nakanomyo, Jutarou Fukazawa, Sayaka Matsui, Yuuki Sakai for technical assistance. This work was supported by JSPS KAKENHI (Grant Number JP22H02676) to Y.H., Takeda Science Foundation to Y.H., GteX Program Japan (JPMJGX23B0) to K.I., the Program for Forming Japan’s Peak Research Universities (J-PEAKS) from JSPS to KI, BBSRC BB/T007117/1 to J.H and BBSRC BB/F011458/1 for confocal microscopy. F.R. is a Leverhulme Early Career Fellow (ECF-2023-534) funded by the Leverhulme Trust and the Isaac Newton Trust (23.08(f)), I.B. is funded by the Herschel Smith Fund studentship.

## Author contributions

G.T. and Y.H. conceived and designed the research. G.T., S.Y., F.R., I.B. and M.S. performed the experiments. G.T., S.Y., F.R., M.S. and Y.H. analyzed the data. K.I. contributed materials and analysis tools. F.R., T.K., J.H. and Y.H. supervised the project. G.T. and Y.H. wrote the manuscript. All authors reviewed and edited the manuscript.

## Declaration of Interests

The authors declare no competing interests.

## Methods

### Plant materials and growth conditions

*Marchantia polymorpha* male Takaragaike-1 (Tak-1) accession was used as wild type in this study. *M. polymorpha* plants were grown on half-strength Gamborg B5 medium (pH 5.5) solidified with 1.4% agar. *M. polymorpha* plants were grown at 22 ℃ under continuous white light. Transgenic plants are listed in Table S1.

### Plasmid Construction

Primers used in this study are listed in Table S2. All plant transformation vectors were generated using the Gateway cloning system (Thermo Fisher Scientific, Waltham, MA, United States). Gateway destination vectors are described in Ishida *et al*. (2022)^32^, Ishizaki *et al*. (2015)^35^ and Sugano *et al*. (2018)^36^.

For genome editing of Mp*GMFR*/Mp*ERF14* and Mp*GCAM1*, guide RNAs were designed using CRISPRdirect^37^. The plasmids for genome editing were constructed according to Sugano *et al*. (2018)^36^.

For the complementation study of genome editing alleles for Mp*GMFR*, a 5026 bp Mp*GMFR* promoter sequence was PCR amplified with a primer pair of MpGMFR_prom_F_InFusion_XbaI and MpGMFR_prom_R_InFusion_XbaI, and cloned into the Xba I digestion site of pMpGWB301 and pMpGWB312 vectors using In-Fusion HD Cloning Kit (Takara Bio, Shiga, Japan) to produce pMpGWB301-proMpGMFR and pMpGWB312-proMpGMFR, respectively. The coding sequence of Mp*GMFR* was PCR amplified from *M. polymorpha* cDNA with a primer pair of MpGMFR_CDS_F and MpGMFR_CDS_R_+stop or MpGMFR_CDS_R_-stop, and cloned into pENTR4 Dual Selection Vector (Thermo Fisher Scientific). To introduce gRNA-resistant mutation to the resulting plasmids, pENTR_MpGMFR_CDS_+stop and pENTR_MpGMFR_CDS_-stop, site directed mutagenesis was performed. The plasmids were PCR amplified with mutagenesis primers, MpGMFR_CDS_gRNAres_F and MpGMFR_CDS_gRNAres_R, and subjected to digestion with Dpn I (Takara Bio), followed by transformation of *Escherichia coli*. Mutagenized plasmids were selected by DNA sequencing. pENTR-MpGMFR_CDS_gRNAres_+stop was transferred to pMpGWB301-_pro_MpGMFR, and pENTR-MpGMFR_CDS_gRNAres_-stop was transferred to pMpGWB312-_pro_MpGMFR using Gateway LR Clonase II Enzyme mix (Thermo Fisher Scientific).

For production of the estrogen-inducible artificial microRNA (amiRNA) lines, an amiRNA target sequence was designed at coding sequence of Mp*GMFR* using amiRNA Design Helper^38^ (https://marchantia.info/tools/amir_helper/) to be inserted into Mp*MIR160* backbone. Cloning into the pMpGWB368^32^ plasmid to generate estrogen-inducible amiRNA construct was performed according to Sakai *et al*. (2024)^17^.

For promoter reporter analysis, a 5026 bp DNA fragment of Mp*GMFR* promoter sequence flanking the translational initiation site was PCR amplified with a primer pair of MpGMFR_prom_F and MpGMFR_prom_R, and cloned into pENTR/D-TOPO vector (Thermo Fisher Scientific). The resulting plasmid, pENTR-proMpGMFR, was transferred to the pMpGWB323-H2B vector^24^ using Gateway LR Clonase II Enzyme mix.

For production of Mp*GMFR* overexpression alleles, pENTR-MpGMFR_CDS_+stop was transferred to pMpGWB303 using Gateway LR Clonase II Enzyme mix. For production of inducible Mp*GMFR* overexpression alleles, pENTR-MpGMFR_CDS_-stop was transferred to pMpGWB113. For the _*pro*_*35S*:Mp*GMFR-mVenus*, the CDS of Mp*GMFR* was synthesized (Genewiz) as a L0_CDS12 part and directly cloned into the pBy12 vector as described in Romani *et al*. (2024)^39^.

For the functional analysis for Mp*ERF1* and Mp*ERF20*/*LAXR*, the coding sequences of Mp*ERF1* and Mp*ERF20*/*LAXR* were PCR amplified from *M. polymorpha* cDNA with a primer pair of MpERF1_CDS_F and MpERF1_CDS_R_-stop or MpLAXR_CDS_F and MpLAXR_CDS_R_-stop, and cloned into pENTR4 Dual Selection Vector. The resulting plasmid, pENTR-MpERF1_CDS_-stop or pENTR-MpLAXR_CDS_-stop were transferred into pMpGWB313 or pMpGWB312-proMpGMFR using Gateway LR Clonase II Enzyme mix.

### Generation of transgenic plants

Agrobacterium-mediated transformation of *M. polymorpha* was performed using regenerating thalli according to Kubota *et al*. (2013)^40^. CRISPR/Cas9-based genome editing was performed according to Sugano *et al*. (2018)^36^. Mutations in the guide RNA target loci were examined by direct sequencing of PCR product amplified from genome DNA samples with primers listed in Table S2.

### Imaging and phenotypic measurement

For the analysis of overall plant morphology and the measurement for the number of gemma cup, plants were imaged under a digital microscope (DMS1000, Leica Microsystems, Wetzlar, Germany) or a digital camera (TG-6, Olympus).

For fluorescence observation in confocal imaging, plants were fixed and cleared with iTOMEI protocol^29^ as described in Takahashi *et al*. (2023)^24^. The cleared samples were mounted in the mounting solution and observed under a confocal laser scanning microscopy (Fluoview FV3000, Olympus). For the observation of developing gemmae and gemma cups, hand-sections were prepared with a scalpel. For 3D-reconstruction, Z-series images were processed using ‘3D-viewer’ or ‘3D-projection’ function of Fiji software^41^.

For the scanning electron microscopy imaging, plants were pre-fixed with 4 % glutaraldehyde in 0.05 M phosphate buffer (pH 7.2) for 2 hours at room temperature, followed by washing in the phosphate buffer. Pre-fixed plants were post-fixed with 1 % osmium tetroxide in 0.05 M phosphate buffer (pH 7.2) for 2 hours at 4 ℃. Fixed plants were dehydrated using ethanol series at room temperature. The plants were immersed in *t*-butyl alcohol and freeze-dried in an evacuator (VFD-21S, Vacuum Device Inc.) until completely dry. Finally, samples were coated with gold by an ion sputtering device (JFC1500, JEOL) and observed under a scanning electron microscopy (JSM-T220A, JEOL).

### GUS staining

GUS staining was performed according to Hirakawa *et al*. (2020)^21^. Briefly, individual plants were stained separately in 30–50 µL GUS staining solution (50 mM sodium phosphate buffer pH7.2, 1 mM potassium-ferrocyanide, 1 mM potassium-ferricyanide, 10 mM EDTA, 0.01 % Triton X-100 and 1mM 5-bromo-4-chloro-3-indolyl-b-D-glucuronic acid) at 37 ℃ in dark. GUS-stained samples were washed with water, cleared with ethanol, and mounted with clearing solution (chloral hydrate-glycerol-water, 8:1:2) for imaging under a light microscope (BX51, Olympus, Tokyo, Japan).

### RT-qPCR

To quantify Mp*GMFR* and Mp*GCAM1* mRNA levels, total RNA was extracted from thallus using NucleoSpin RNA Plant (Macherey-Nagel, Duren, Germany) according to manufacturer’s instruction. First-strand cDNAs were prepared using ReverTra Ace qPCR RT Master Mix with gDNA Remover (TOYOBO, Osaka, Japan). RT-qPCR was performed with the reagents, TB Green Premix Ex Taq II (Takara Bio) and THUNDERBIRD Next SYBR qPCR Mix (TOYOBO), using the devices, Step One Plus Real-Time PCR System (Thermo Fisher Scientific) and CFX Connect (Bio-Rad Laboratories, California, USA). Mp*APT* (Mp*3g25140*) was used as a reference gene.

### Phylogenetic analysis

Protein sequences were retrieved from the following databases: MarpolBase^42^ (https://marchantia.info), Phytozome^43^ (https://phytozome-next.jgi.doe.gov/), TAIR (http://www.arabidopsis.org/) and GinkgoDB^44^ (https://ginkgo.zju.edu.cn/genome/). Alignment was performed on the amino acid sequences of the AP2 domain using CLUSTALW (https://www.genome.jp/tools-bin/clustalw). After manually removing the alignment gaps using SeaView^45^, phylogenetic analysis was performed on the alignment using MrBayes3.2.7^46^. Two runs with four chains of Markov chain Monte Carlo (MCMC) iterations were performed for 6,000,000 generations, keeping one tree every 100 generations. The first 25% of the generations were discarded as burn-in and the remaining trees were used to calculate a 50% majority-rule tree. The standard deviation for the two MCMC iteration runs was below 0.01, suggesting that it was sufficient for the convergence of the two runs. Convergence was assessed by visual inspection of the plot of the log likelihood scores of the two runs calculated by MrBayes^47^. Character matrix used to run the Bayesian phylogenetic analysis is provided in Data S1.

### Data visualization

The statistical software R version 4.5.0 was used for data visualization.

### Quantification and statistical analysis

For phenotypic quantification, JMP pro 18 (JMP Statistical Discovery LLC, North Carolina, USA) was used for statistical tests. Statistical details including the type of test, sample size and statistical significance can be found in figure legends.

## Notes

### Competing Interest Statement

The authors have declared no competing interest.

