## Supplemental Figures for "Initiation of asexual reproduction by the AP2/ERF gene *GEMMIFER* in *Marchantia polymorpha*"

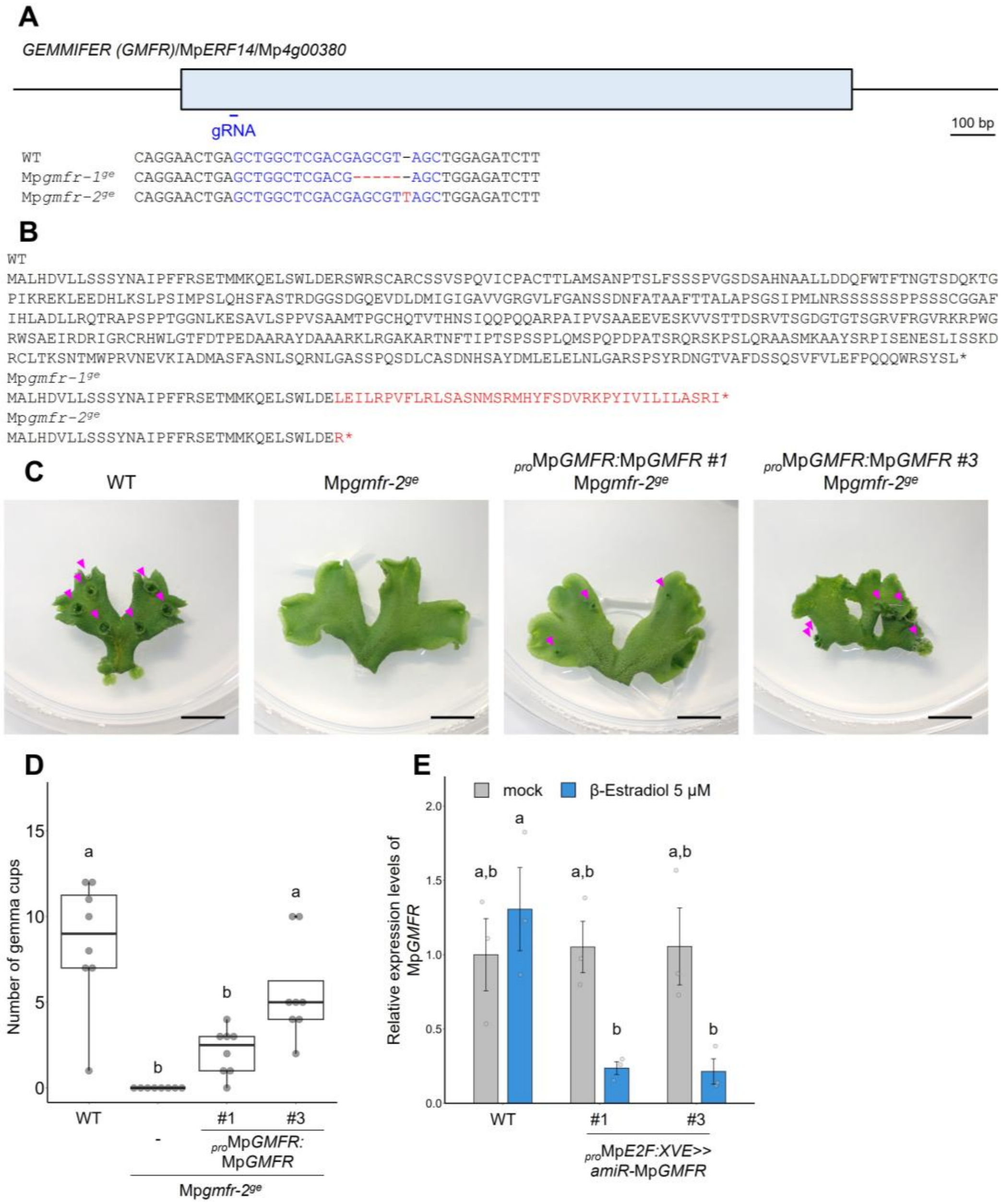


**Fig. S1 | Loss-of-function mutants for *GEMMIFER* (Mp*GMFR*/Mp*ERF14*)**
(**A**) Structure of Mp*GMFR*/Mp*ERF14* locus with the position of designed guide RNA (gRNA). Exons are shown as boxes. Genotyping of genome editing alleles are indicated below. gRNA sequence is in blue. Deleted or inserted bases are indicated in red. (**B**) WT and mutant protein sequences deduced from the genomic DNA sequences are indicated. Sequences different from WT are in red. Asterisks indicate translational termination. (**C**) Complementation test for the Mp*gmfr*$^{ge}$ allele by gRNA-resistant Mp*GMFR* coding sequence driven under own promoter. Magenta arrowheads indicate gemma cups. (**D**) The number of gemma cups (n=8). (**E**) Relative expression levels of Mp*GMFR* in 14-day-old plants of wild type and $_{pro}$Mp*E2F:XVE*>>*amiR*-Mp*GMFR* grown from gemmae, normalized by Mp*APT* (n=3).
In **D,** the boxes show the median and interquartile range, and the whiskers show the 1.5× interquartile range. Individual data points are plotted as dots. In **E,** data are represented by mean

(bars) and individual data points (dots). Two-way ANOVA with Tukey's post hoc test in **D,E**. Means sharing the superscripts are not significantly different from each other, $p < 0.05$. Scale bars represent 1 cm in **C.**

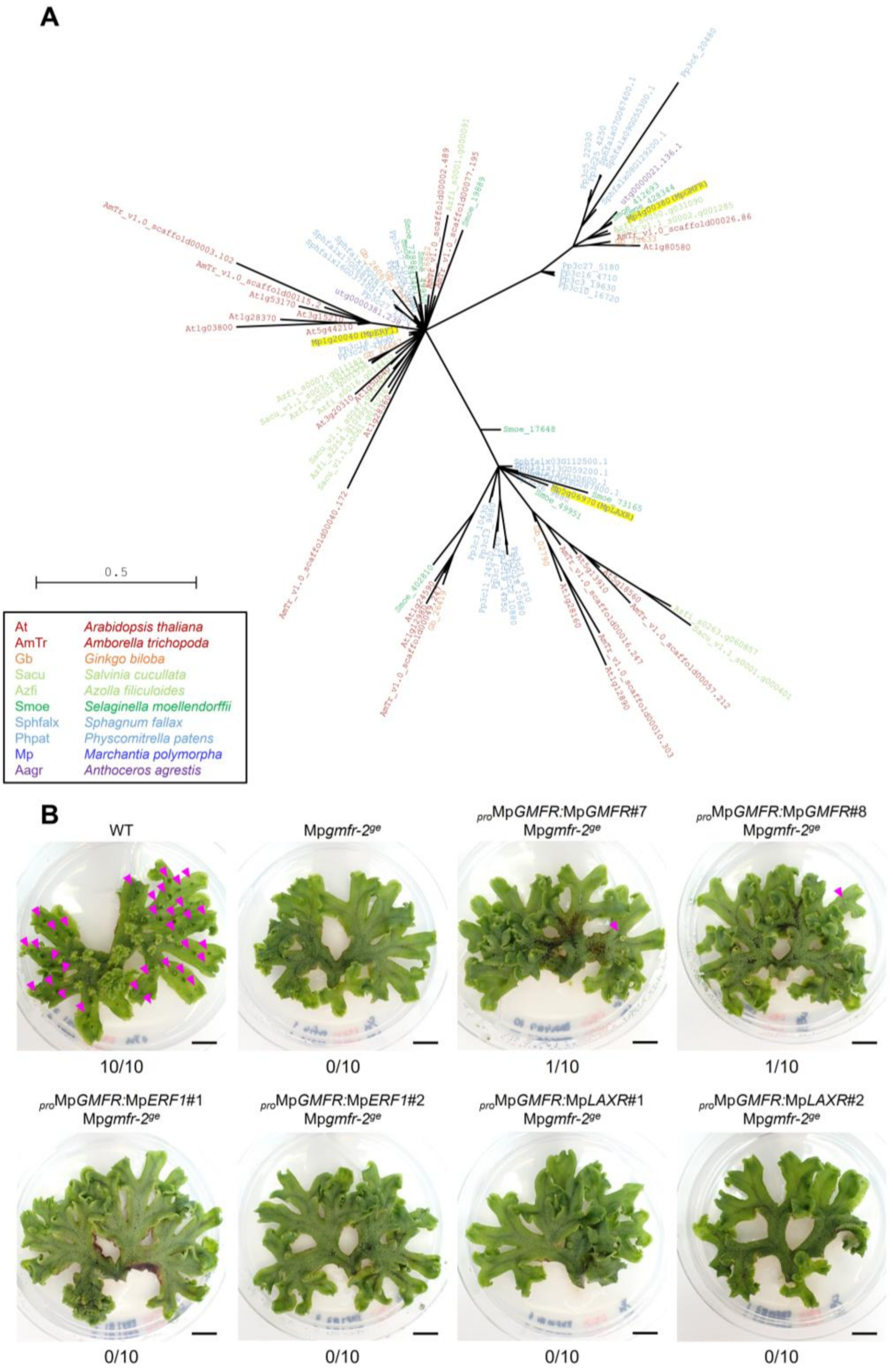


**Fig. S2 | Analysis of ERF-VIII genes**

(**A**) Molecular phylogenetic tree of class VIII of AP2/ERF family among land plant species, generated with a Bayesian method based on the conserved AP2 domain. (**B**) Complementation test of Mp*gmfr*$^{ge}$ by Mp*GMFR-GR*, Mp*ERF1-GR* and Mp*LAXR-GR* driven under Mp*GMFR* promoter. The ratio of thallus containing at least one gemma cup is indicated below the panels (n=10). Magenta arrowheads indicate gemma cups. Scale bars represent 1 cm.

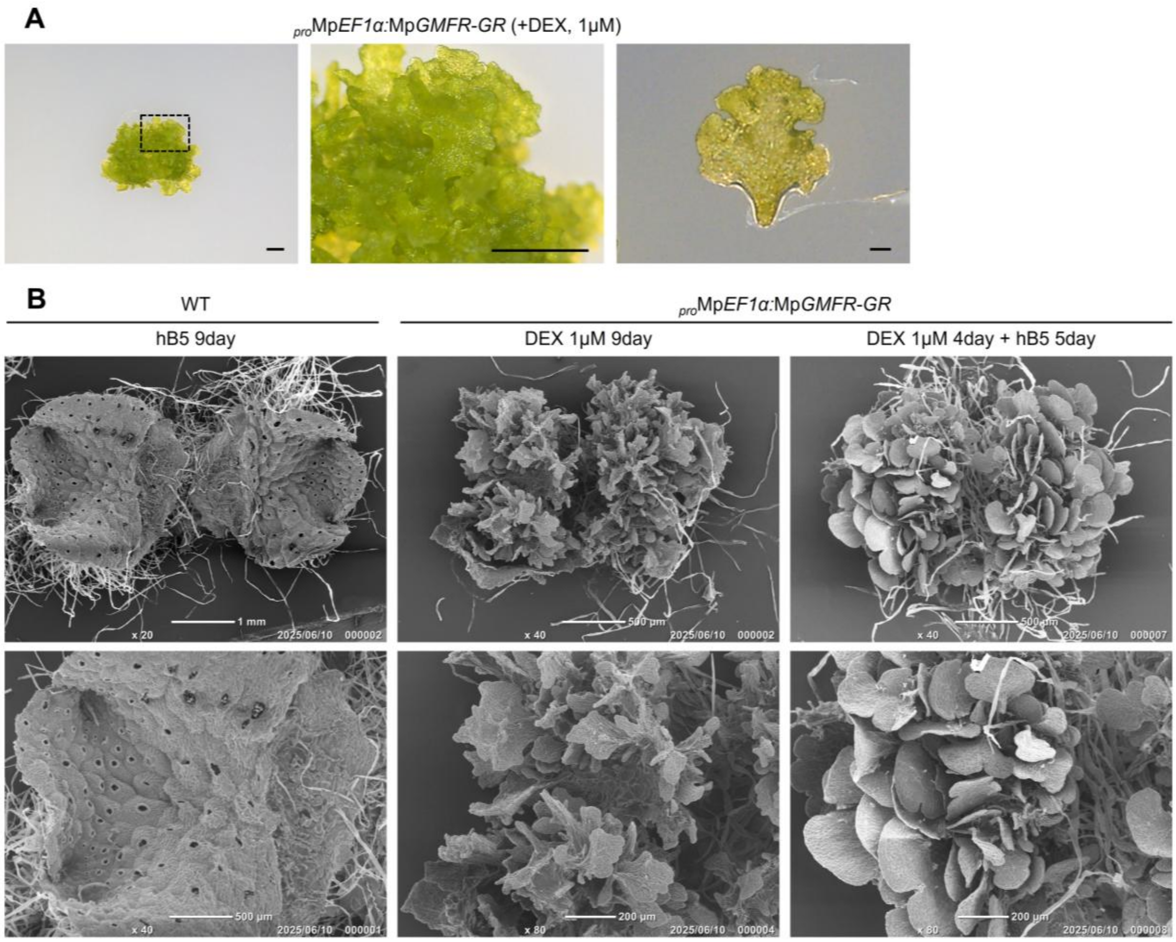


**Fig. S3 | Continuous overexpression of Mp*GMFR* forms immature small thalli**

(**A**) Morphology of $_{pro}$Mp*EF1α:*Mp*GMFR-GR* plant grown for 9 days on 1 μM DEX-containing medium. Scale bars represent 1 mm (left and center) or 100 μm (right). (**B**) SEM images of the surface of 9-day-old thalli. Genotypes and growth conditions are indicated above the panels.

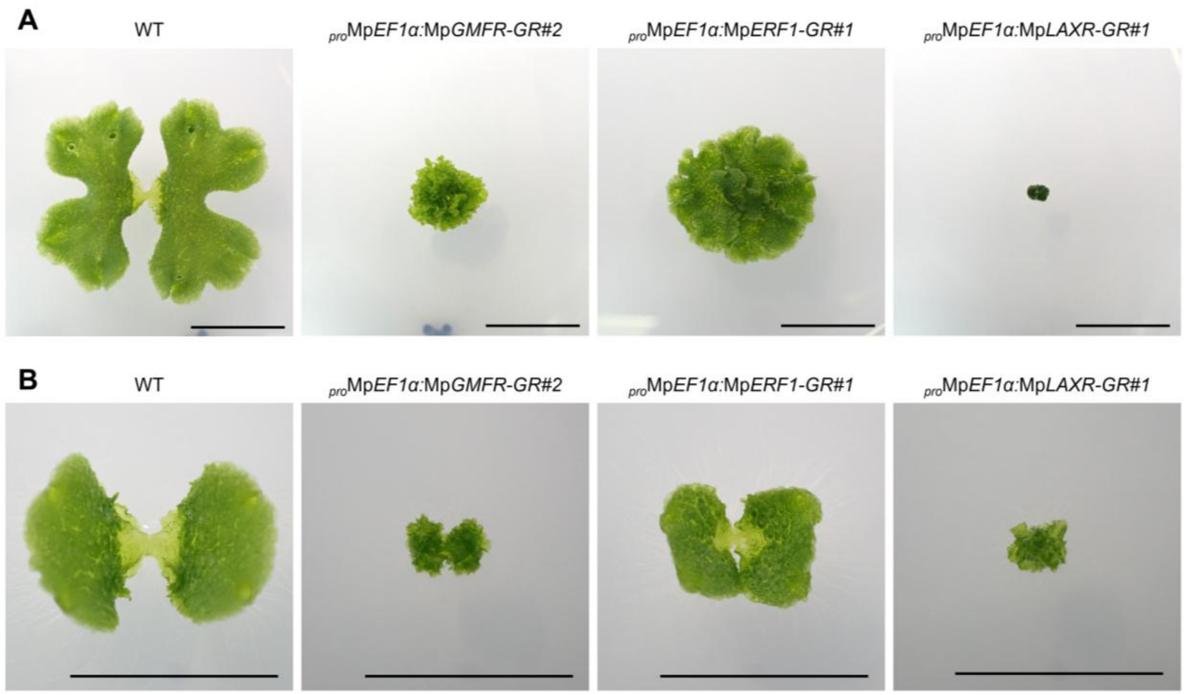


**Fig. S4 | Overexpression phenotypes of Mp*ERF1* and Mp*LAXR*/Mp*ERF20***

(**A–B**) Phenotypes of inducible overexpression of Mp*GMFR*, Mp*ERF1* and Mp*LAXR*/Mp*ERF20*. Overall morphology of plants grown on 1 μM DEX-containing medium for 9 days (**A**) and those grown on 1 μM DEX-containing medium for 4 days followed by culture on DEX-free medium for 5 days (**B**). Scale bars represent 1 cm in **A** and **B**.

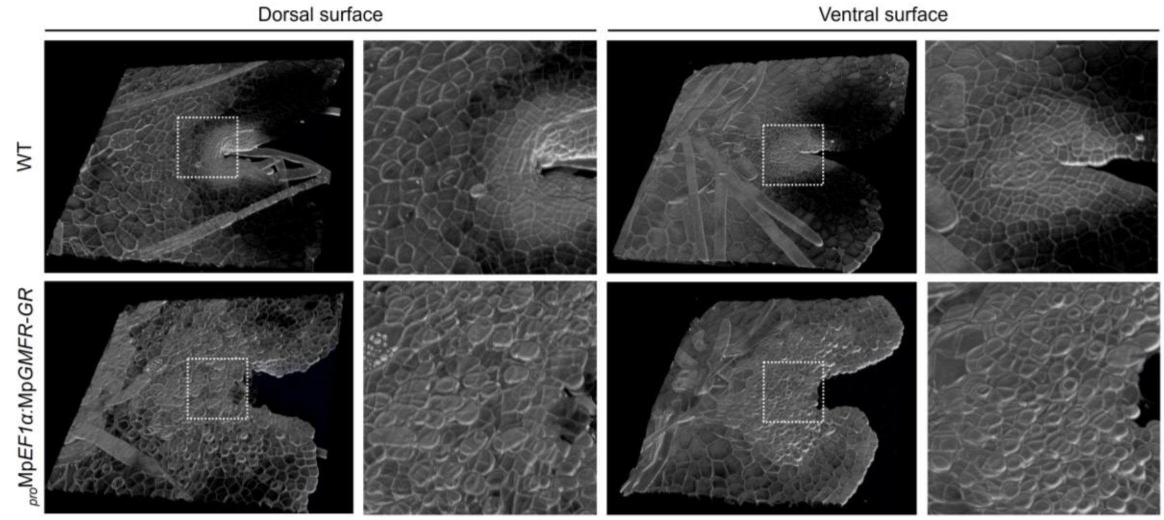


**Fig. S5 | MpGMFR-induced protruding cells are formed on both dorsal and ventral surfaces**

3D-reconstructed images of the 3-day-old apical notch in wild-type and $_{pro}$Mp*EF1α:*Mp*GMFR-GR*. The right panel shows the magnified image of a dashed line square in the left panel. Cell walls were stained with SR2200.

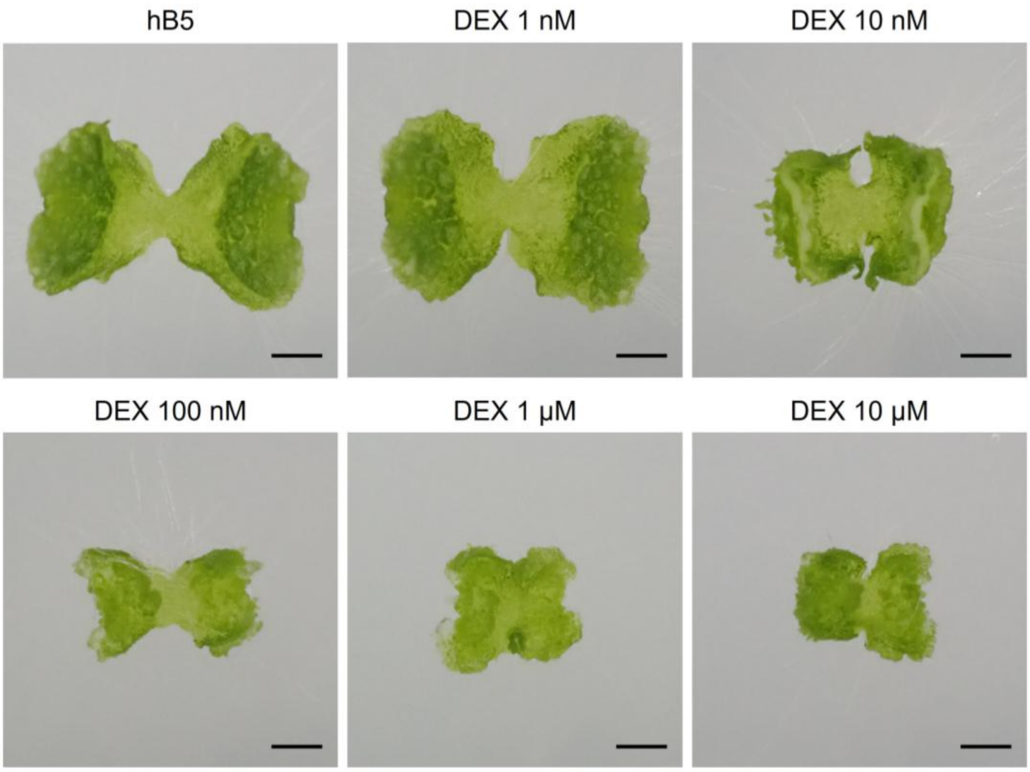


**Fig. S6 | Effects of various concentrations of DEX in $_{pro}$Mp*EF1α:*Mp*GMFR-GR* plants**

Effects of various concentrations of DEX in 7-day-old $_{pro}$Mp*EF1α:*Mp*GMFR-GR* plants grown from gemmae. Growth conditions are indicated above each panel. Scale bars represent 1 mm.

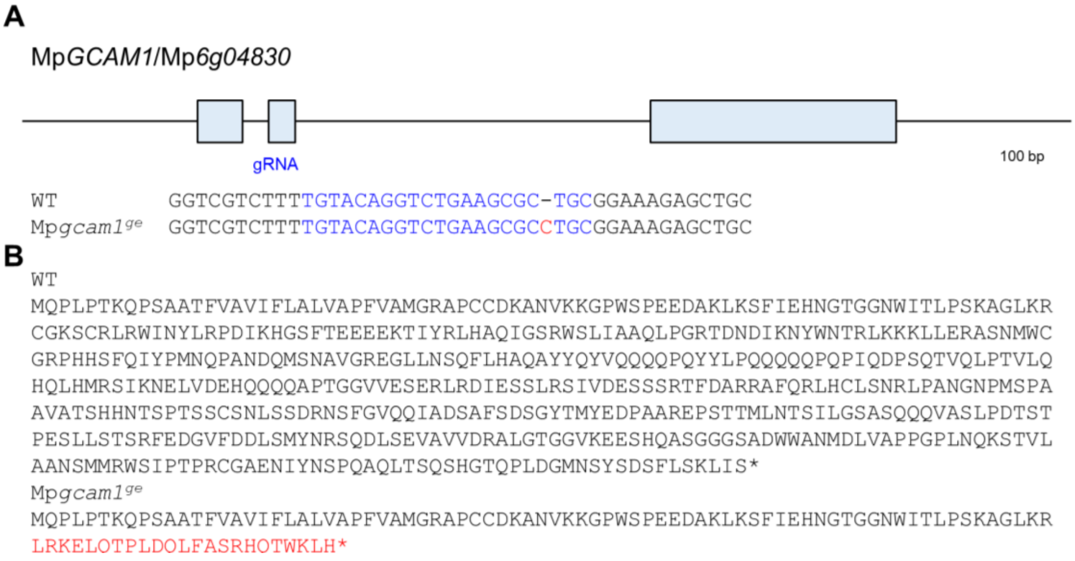


**Fig. S7 | Loss-of-function mutants for Mp*GCAM1***

(**A**) Structure of Mp*GCAM1*/Mp6g04830 locus with the position of designed guide RNA (gRNA). Exons are shown as boxes. Genotyping of genome editing alleles are indicated below. gRNA sequence is in blue. Deleted or inserted bases are indicated in red. (**B**) WT and mutant protein sequences deduced from the genomic DNA sequences are indicated. Sequences different from WT are in red. Asterisks indicate translational termination.

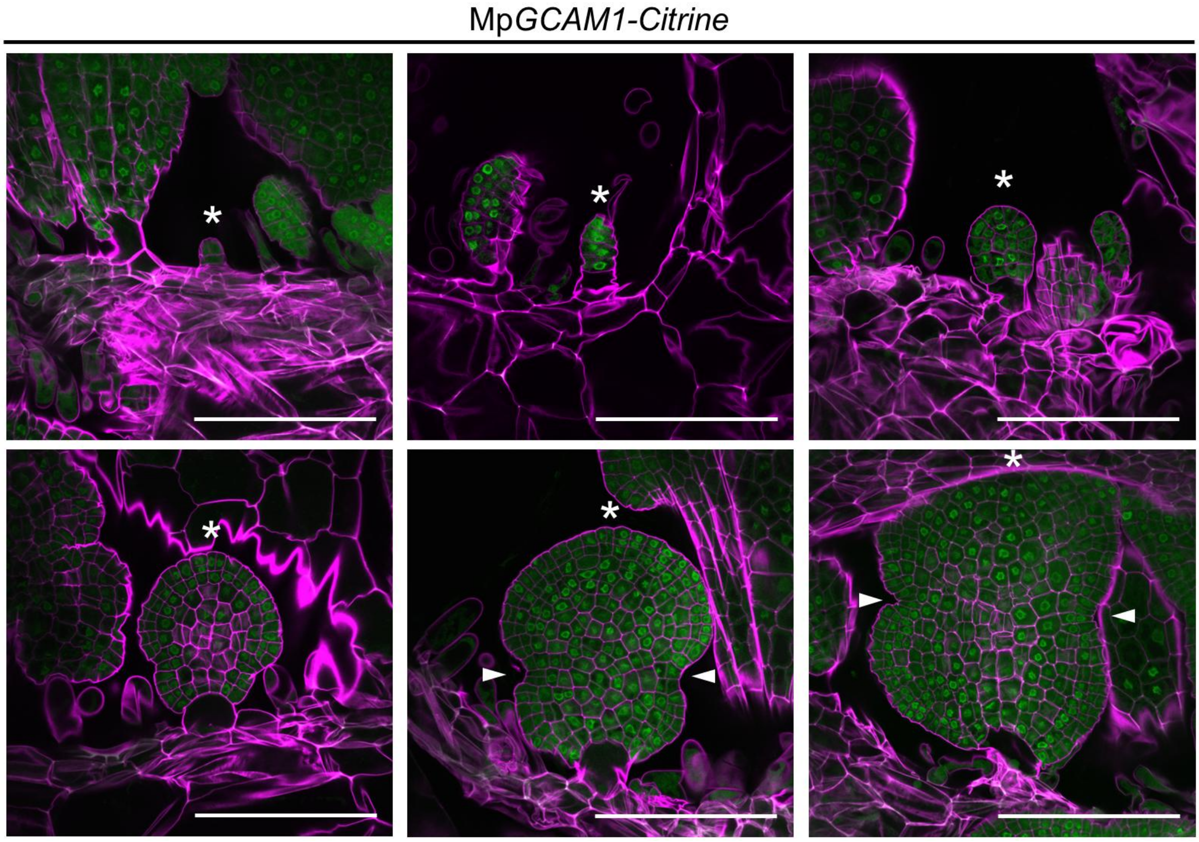


**Fig. S8 | Expression patterns of Mp*GCAM1* during the development of gemmae**

Confocal imaging of Mp*GCAM1-Citrine* plants in gemma development. Asterisks indicate developing gemmae. Arrowheads indicate apical notches. Cell walls were stained with SR2200. Scale bars represent 100 µm.
